# The allosteric mechanism of substrate-specific transport in SLC6 is mediated by a volumetric sensor

**DOI:** 10.1101/555565

**Authors:** Michael V. LeVine, Daniel S. Terry, George Khelashvili, Zarek S. Siegel, Matthias Quick, Jonathan A. Javitch, Scott C. Blanchard, Harel Weinstein

## Abstract

Neurotransmitter:sodium symporters (NSS) in the SLC6 family terminate neurotransmission by coupling the thermodynamically favorable transport of ions to the thermodynamically unfavorable transport of neurotransmitter back into presynaptic neurons. While a combination of structural, functional, and computational studies on LeuT, a bacterial NSS homolog, has provided critical insight into the mechanism of sodium-coupled transport, the mechanism underlying substrate-specific transport rates is still not understood. We present a combination of MD simulations, single-molecule FRET imaging, and measurements of Na^+^ binding and substrate transport that reveal an allosteric mechanism in which residues F259 and I359 in the substrate binding pocket couple substrate binding to Na^+^ release from the Na2 site through allosteric modulation of the stability of a partially-open, inward-facing state. We propose a new model for transport selectivity in which the two residues act as a volumetric sensor that inhibits the transport of bulky amino acids.

## Introduction

Neurotransmitter:sodium symporters (NSS) in the SLC6 family are secondary active transport proteins that regulate the extent and duration of neurotransmission events by re-importing neurotransmitter molecules following their release into the post-synaptic space (1). Highlighting their critical function in regulating neurotransmission, small molecules targeting these transporters include clinically important antidepressants and psychostimulant drugs of abuse (2, 3). The transport mechanism of NSS family members is thought to involve conformational changes that control accessibility of the transporter to the intra- and extra-cellular milieu of the cell in alternating fashion (4), with net transport of the substrate against its concentration gradient powered by the transmembrane electrochemical Na^+^ gradient.

While the fundamental thermodynamics of secondary active transport are well understood, intense study is currently focused on the identification of the specific sequence of substrate binding and release events, as well as the series of conformational changes that are coupled to and facilitate those processes (5, 6, 15–17, 7–14). X-ray crystallographic studies have provided insight into the NSS molecular architecture, beginning with crystal structures of the Na^+^-coupled prokaryotic amino acid transporter LeuT (18, 19) that allowed for existing mechanistic models to be put into a structural context (20). Single-molecule FRET (smFRET) studies have illuminated the dynamic sampling of functionally important intermediate states visited by the transporter during the transport cycle (21–23). Computational studies employing Molecular Dynamics (MD) simulation have identified key elements of the complex allosteric mechanism by which conformational changes in the transporter are coupled to the binding and release of ions and substrates (1, 14, 17, 24–30).

Our previously described smFRET experiments (21–23) reported on the dynamics and displacement of the N-terminus from its central position in the intracellular gate. This critical network includes a salt bridge and hydrogen bond interactions that maintain an inward-closed conformation in which the substrate is occluded from the intracellular solvent. These results were obtained by measuring energy transfer between fluorophores attached through thiol-specific chemistry to amino acid position 7 in the N-terminus and position 86 in intracellular loop 2 (IL2). We initially reported two states: a high-FRET state that likely corresponds to an inward-closed conformation (denoted IC) and a low-FRET state that likely corresponds to an inward-open conformation (denoted IO) in which the N-terminus has dissociated from the intracellular gate such that the substrate is exposed to the intracellular solvent. These studies (21–23), and later experiments utilizing electron pair resonance (EPR)(31) and cysteine accessibility (29), probed the modulation of the IO/IC exchange process by substrate binding in the primary substrate (S1) binding site and in the allosteric secondary substrate (S2) binding site.

Leveraging technical advances in the smFRET technology, including sCMOS imaging (32) and self-healing fluorophores (33, 34), we recently identified in LeuT an intermediate-FRET state that was shown to be consistent with a partially inward-open conformation (23) (denoted here as IO^2^). This state was rapidly sampled in the presence of Na^+^ and the substrate Ala. However, replacing Ala with the slowly transported substrate Leu, or replacing Na^+^ with Li^+^, which doesn’t support transport, stabilized the IC state and slowed the overall dynamics. In addition, we found that the Ala-stabilized IO^2^ state was selectively destabilized by high concentrations of Na^+^, suggesting that this state corresponds to a conformation in which only one Na^+^ binding site is occupied, while the other Na^+^ has been released. Computational (14, 17, 35) and experimental (17) evidence suggests that it is the Na^+^ in the Na2 site that is released, a process that is thought to be the obligatory first step of the substrate release mechanism. As Ala demonstrates a substantially greater maximum velocity of transport (*V_max_*) than Leu, we hypothesized that substrate-specific stabilization of the IO^2^ state may trigger release of Na^+^ from the Na2 site and allow for transport to proceed, whereas substrate-specific stabilization of the IC state may inhibit Na^+^ release to the extent to which it becomes rate-determining. This model is consistent with thermodynamic coupling function (TCF) (36) and Markov State Model analyses performed for the human dopamine transporter, which revealed an allosteric coupling between Na^+^ release from the Na2 site and inward-opening (13, 14, 35).

Despite these advances in quantifying substrate-specific allosteric modulation of intracellular gating and Na^+^ release by substrates, the phenomenon is still not understood at a molecular level. In order to obtain such detailed mechanistic information, we (14, 24, 36, 37) and others (38–41), have developed several computational methods and theoretical models, including N-body Information Theory (NbIT) analysis (24, 37) and the TCF theory of allostery (14, 36). Previously, we applied NbIT analysis to Molecular Dynamics (MD) simulation of LeuT bound to Na^+^ and Leu in the occluded state (24) and were able to identify significant information transmission between the S1 site and the intracellular gate mediated by transmembrane helix (TM) 6b through the direct interaction of substrate with the side chain of the conserved F259 (and to a lesser extent by TM8 and I359). Interestingly, a comparative analysis of crystal structures of LeuT bound to a number of substrates with different rates of transport (including Gly, Ala, Leu, Met, L-4-fluorophenylalanine (L-4-FP), and Trp) revealed characteristic differences in their mode of interaction with F259 (19). Specifically, Trp (like L-4-FP) is seen to makes strong ring-ring contacts with F259, and measurements show that it is not transported but rather acts as an inhibitor. Met and Leu have weaker hydrophobic interactions with F259, and are slowly transported substrates; whereas Ala makes no contact with F259 and is an efficiently transported substrate. A similar (albeit smaller) difference in the mode of interaction of I359 with the substrates is also apparent. This set of relations between the extent of interaction with F259 and the differences in transport efficacy, combined with the substrate-specific allosteric modulation observed for Leu and Ala in our previous smFRET experiments (22, 23), led us to hypothesize a mechanistic relationship (1). According to this hypothesis, substrates that interact strongly with F259 stabilize the IC state and quench the gating dynamics required for transport, whereas substrates that interact weakly with F259 stabilize the IO^2^ state, trigger release from of Na^+^ from the Na2 site and facilitate transport.

Here we have tested the mechanistic hypothesis that substrate-induced inward closing is mediated by the extent of substrate interaction with F259 and neighboring I359. To this end, we combined microsecond-scale MD simulations, hybrid quantum mechanics/molecular mechanics (QM/MM) calculations, and transport and smFRET experiments in the analysis of LeuT in complex with the substrates Gly, Ala, Val, and Leu. Our results suggest a mechanism in which F259, assisted by I359, acts as a volumetric sensor that allosterically couples the chemical identity of the substrate to different extents of inward closing in a manner that substrate-specifically modulates the intracellular release of Na^+^ from the Na2 site.

## Results

### Differential modulation of F259 and I359 rotamer dynamics by substrates

LeuT can bind and transport a number of amino acid substrates (19). Whereas the transport and binding kinetics vary significantly among these different substrates (including Gly, Ala, Val and Leu), the crystal structures of LeuT in complex with each of these ligands displayed only subtle changes (19). Specifically, the Ala- and Gly-bound structures display only a 30**°** rotation of the F259 χ_2_ angle and an ~15° rotation of the I359 χ_1_ angle when compared to the Leu-bound structure. Because we had previously identified both of these residues as significant mediators of information transmission between the S1 binding site and the intracellular gate in the Leu-bound state(24), we hypothesized that there exists a relation between substrate-specific differences in experimentally measured *V_max_* and the modulation of the conformational dynamics of F259 and I359. To investigate the possibility that subtle rotations of F259 and I359 identified in the X-ray structure reflect larger differences in conformational flexibility or heterogeneity under physiological conditions, we turned to a panel of protein-substrate complexes that allowed for a systematic evaluation of how the graded reduction of the interaction between substrate and the F259 and I359 side chains may affect the conformational dynamics of these two residues.

Microsecond-scale all-atom MD simulations and rotamer scan calculations with the combined quantum mechanical/molecular mechanical approach (QM/MM), were carried out for LeuT in complex with Gly, Ala, Val, and Leu (see **Methods**). The results showed that both F259 and I359 sample multiple conformational states at room temperature, and do so in a substrate-dependent manner. In particular, we found that when LeuT is bound to Ala and Gly, which have relatively small side chains, F259 freely rotates between four χ_2_ angle states (**Fig. 1A-B; Figs. S1, S2, S7**). The χ_2_ angle describes the rotation of the phenyl ring relative to the Cα-Cβ bond. Due to the symmetric nature of the phenyl ring, these four rotamer states are composed of two sets of two symmetric rotameric states, referred to here as “perpendicular” and “parallel”. In the perpendicular state, the F259 phenyl side chain faces the substrate (**Fig. 1A,** χ_2_ angles of approximately ~75**°** and ~255**°**), whereas in the parallel state, the side chain turns approximately 90**°** (χ_2_ angles of approximately ~ −5**°** and ~165**°**) and is aligned with the axis of the substrate. The free exchange between perpendicular and parallel states observed in our simulations is consistent with the behavior of an ideal allosteric channel in the allosteric Ising model (42).

**Figure 1.**
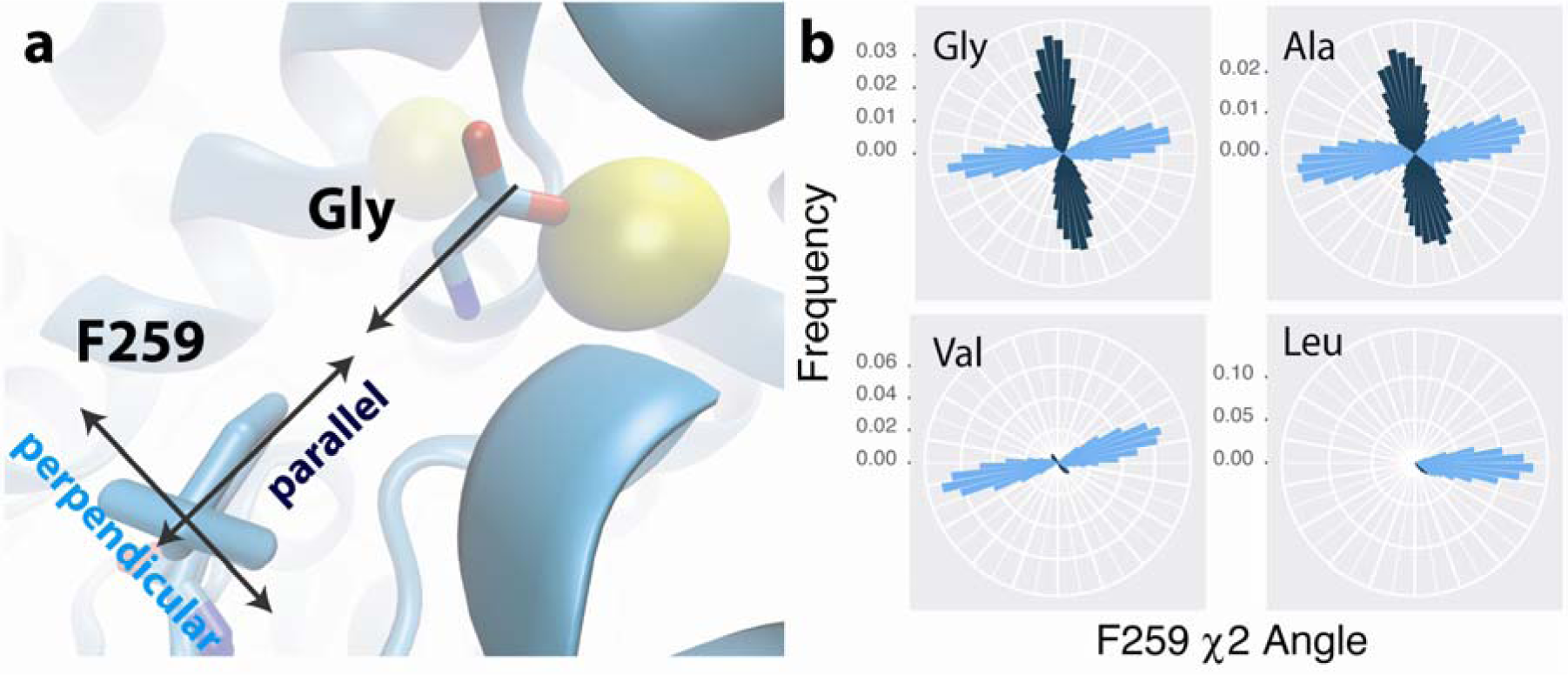
Substrates differentially modulate F259 rotameric dynamics. (**A**) The two possible states of the F259 side chain. The substrate glycine is shown for reference, with surrounding Na1 (right) and Na2 (left) shown in yellow. (**B**) Rose plots showing the angular histograms of the F259 χ_2_ angle sampled during the converged portion of the MD simulations, colored by angular k-means clustering.

When LeuT is bound to Val, a substrate with a larger side chain, the F259 side chain is stabilized in the perpendicular state (**Fig. 1B; Fig. S3**), as observed in the X-ray crystal structures (19), but still exhibits free rotation between the symmetric perpendicular states on the nanosecond time scale. Interestingly, the dynamics reveal that while F259 freely rotates in both directions in the presence of Gly and Ala, the ring shows a preference for clockwise motion in the presence of Val (see. **Fig. S7**). The Leu-bound complex shows only extremely rare exchange between the symmetric sub-states of the F259 rotamers (**Fig. 1B; Fig. S4, S7**; see **Methods** for details of convergence analysis).

We also observe that the I359 side chain can sample two χ_1_ states (**Fig. S6**), which are differentially modulated by the four substrates in a similar fashion (**Figs. S8, S9, S1-4**). The χ_1_ angle describes the rotation of the Ile side chain relative to the Cα-N bond of the backbone. When Leu or Val is bound, I359 predominantly samples a state at 240°. By contrast, I359 predominately samples a state at 120° in the Gly- and Ala-bound simulations, which was not observed in the X-ray structures of these protein:substrate complexes.

The results above show that in the presence of Ala and Gly, the F259 parallel and perpendicular states were equally probable (and I359 spent nearly all of the trajectory in the 120° state). Notably, the empirical rotamer distribution of Phe found in protein structures in the PDB (43) indicates that this rotamer state is not expected to be high probability. Thus, we hypothesized that when F259 is unhindered by the substrate side chain, the parallel state is still enthalpically unstable relative to the perpendicular state, and would thus be masked in a low temperature x-ray crystal structure. In order to investigate this possibility, we scanned the angle space of both F259 and I359 using QM/MM in complexes in which the protein backbone was constrained to the conformation observed in the X-ray structures. We found distinct energy wells in the F259 χ_2_ potential energy landscape in our Gly- and Ala-bound QM/MM calculations (**Fig. S10**) and observe that the parallel state is approximately 3 kcal higher in energy than the perpendicular state that is observed in the X-ray structure, indicating an enthalpic preference consistent with our hypothesis. Surprisingly, however, the parallel minima are not present in the Leu-bound complex, and are roughly 6 kcal higher in energy in the Val-bound state. Similarly, there is a second minimum in the I359 χ_1_ landscape corresponding to the alternative conformation observed in our MD simulations (**Fig. S11**), destabilized by roughly 3 kcal relative to the rotameric state found in the X-ray structure. Again, no alternative state for the I359 χ_1_ angle is present in the Leu-bound complex, and the state is roughly 8 kcal higher in energy in the Val-bound complex. These calculations provide independent support for the substrate-specific modulation of the rotameric state of the S1 binding residues F259 and I359.

### Free rotation of F259 is associated with stabilization of the IO^2^ state

Because our previous findings implicated F259 and I359 in allosteric modulation of intracellular gate dynamics (24), we next sought to identify the relationship between substrate-dependent rotamer dynamics and the observed differences in transport rates for the various substrates. Based on our previous observation (22, 23) that Ala stabilized the inward-facing, intermediate-FRET state whereas Leu did not, we hypothesized that the intermediate-FRET state may be allosterically coupled to the F259 and I359 rotamer equilibriums, and in particular to transitions to the parallel state of F259. Based on this hypothesis and our MD and QM/MM results, we expected that Gly (like Ala) would stabilize the inward-facing, intermediate-FRET state where Val (like Leu) would not.

To test this hypothesis, we performed smFRET imaging of the intracellular dynamics of LeuT (see **Methods** for details) in the presence of Gly, Ala, Val, and Leu and varying Na^+^ concentrations. Ala and Gly both elicited a Na^+^-dependent stabilization of the inward-facing, intermediate-FRET state (**Fig. 2A**) and an increased rate of intracellular gating dynamics (**Fig. 2B,C**) compared to Na^+^ alone (**Fig. S12**). By contrast, Val partially stabilized the inward-closed state and only slightly increased the rate of dynamics, and Leu stabilized the inward-closed state even further and did not increase dynamics compared to Na^+^ alone (**Fig. 2A,B,C**). These results establish a clear correlation between the substrate-modulated F259 rotamer dynamics observed in our MD simulations and QM/MM calculations and the substrate-modulated intracellular gating dynamics measured by smFRET. Importantly, both F259 dynamics and intermediate-FRET state occupancy also follow the same rank order as the rates of transport (*V_max_*) of these substrates (**Table S1**). These findings support our hypothesis that F259 dynamics are allosterically coupled to intracellular gating dynamics.

**Figure 2.**
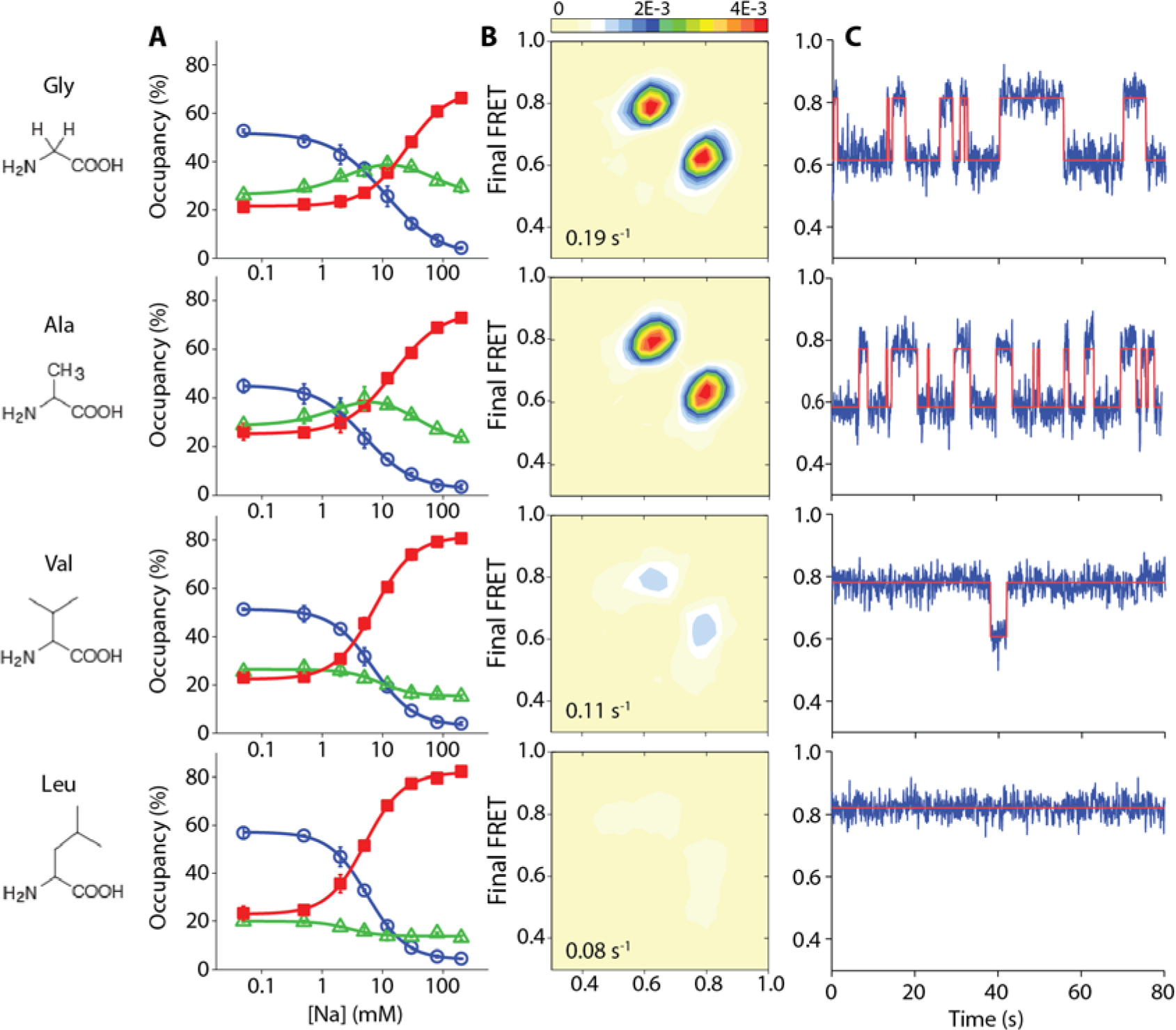
Small substrates stabilize the intermediate state and induce Na^+^ release. (**A**) Intracellular labeled, LeuT-7C/86C was imaged using smFRET in the presence of 10 mM glycine (top row), 300 μM alanine (upper middle), 100 μM valine (lower middle), and 10 μM leucine (bottom row) and the indicated concentrations of Na^+^. Shown are ensemble averaged occupancies in the low-(blue circles), intermediate-(green triangles), and high-FRET (red squares) states. Lines are fits to dose response functions. Error bars are mean ± s.d. of two repeats. (**B**) Transition density contour plots, which show the mean FRET value in the dwell just prior to (bottom axis) and just after (left axis) each transition between distinct FRET states, summed into a histogram for each substrate in the presence of 30 mM Na^+^. The average transition rate is given at the bottom left of each panel. Scale bar at top in transitions per histogram bin per second. (**C**) Representative FRET (blue) and state assignment (red) traces corresponding to the experiments in panel B.

### Mutations at position 259 that favor the parallel rotamer also stabilize the IO^2^ state

We next sought to test the association between F259 rotamer dynamics and the intracellular gating observed in our smFRET experiments by constraining the rotameric state in a substrate-independent manner. Noting that the residue corresponding to F259 is a Trp in the homologous Gly transporter GlyT and that our past homology modeling of GlyT bound to Gly positioned the tryptophan ring in the parallel state (44), we investigated the effect of a F259W mutation in LeuT. We reasoned that because substrate size affects the rotamer dynamics of F259, placing a larger residue (F259W) in the substrate binding site at this position would stabilize the residue in the parallel rotameric state. We first tested this model with a 3 microsecond MD simulation of the LeuT-F259W- in complex with Gly (**Fig. 3A**). In so doing, we found that W259 is indeed locked in the parallel state for the entire trajectory (**Fig. 3B, Fig. S13**), while I359 predominantly occupies the 120° state and displays unimpeded transitions (**Fig. S13**). Consistent with the hypothesized allosteric coupling between rotamer dynamics and intracellular gating, smFRET imaging of the F259W construct showed a stabilization of the intermediate-FRET state (IO^2^), a destabilization of the inward-closed state, and an increase in the apparent rate of dynamics (**Fig. 3C,D**). These effects, which correspond to those induced by Gly and Ala in LeuT-wild-type (WT), occurred in LeuT-F259W even in the absence of substrates. This observation is consistent with the association between occupancy of the parallel rotameric state and occupancy of the IO^2^ state. Notably, dwells in the low- and intermediate-FRET states were more distinct than in LeuT-WT, where these dynamics were difficult to discern due to time averaging at the timescale of imaging (100 ms).

**Figure 3.**
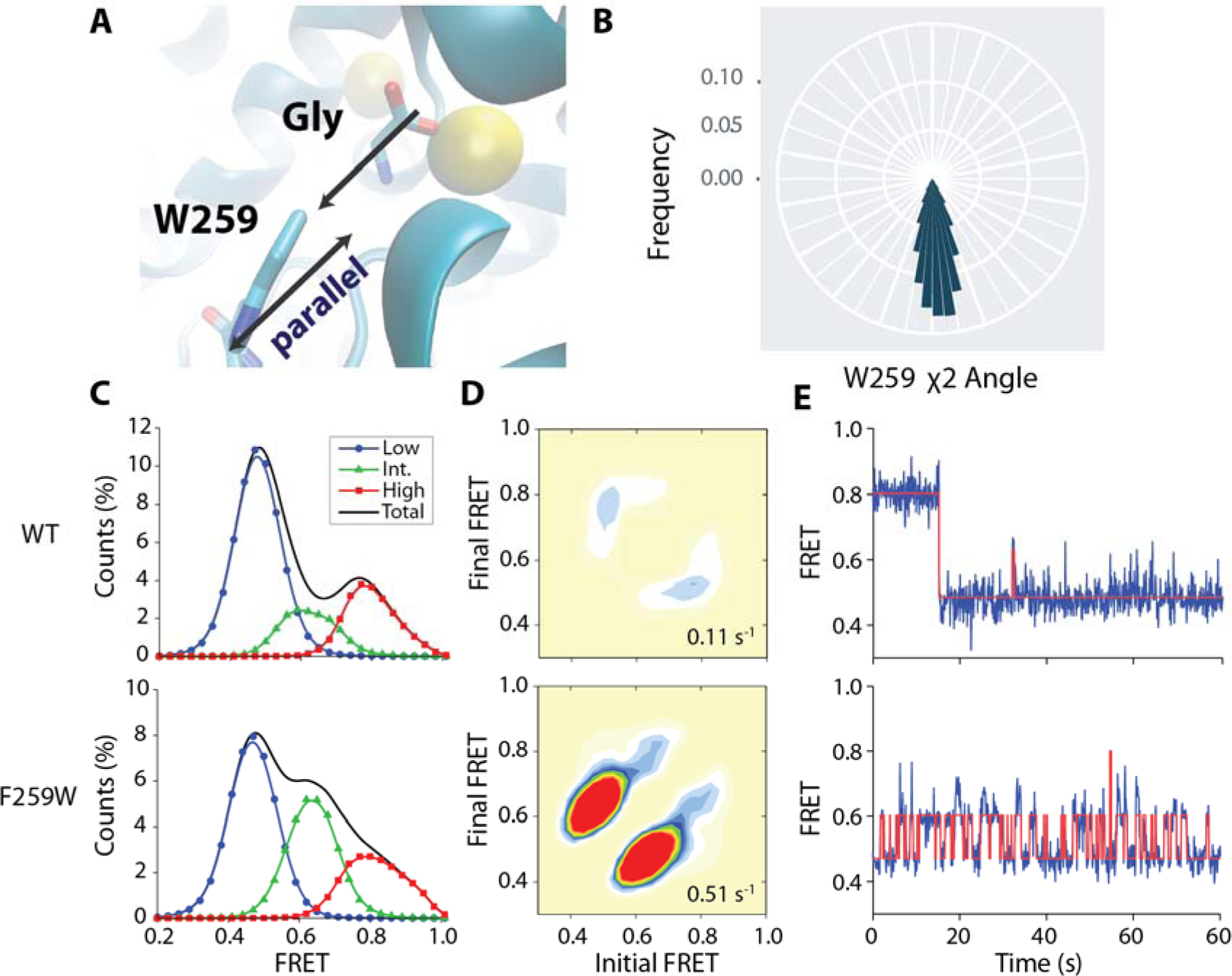
The F259W mutation stabilizes the intermediate FRET state. **(A)** A representative frame of F259W from the MD simulation (depicted as in Fig. 1A). **(B)** Rose plot of the angular histogram of the F259 χ_2_ angle, colored by angular k-means clustering. (**C,D**) LeuT-WT (top panels) and -F259W (bottom panels) were imaged in the absence of Na^+^ and substrates. **C.** FRET histograms for the low-(blue circles), intermediate-(green triangles), and high-FRET (red squares) states and total histogram (black line). (**D**) Transition density contour plots, with the average transition rate in the lower-right corner. (**E**) Example FRET (blue) and state assignment (red) traces.

To test whether the mutation of F259 affects the interaction of the transporter with Na^+^, we measured ^22^Na^+^ binding in the absence of substrate with the scintillation proximity assay. Whereas LeuT-WT exhibited a 2 Na^+^:1 LeuT binding stoichiometry, only one Na^+^ molecule binds to the F259W mutant (**Fig. 4A**). smFRET measurements of Na^+^-induced inward closing of the F259W mutant in the absence of substrates also demonstrated significantly lower affinity for Na^+^ (**Fig. 4B**). That Na^+^ can have a very low affinity for the Na2 site under these conditions is consistent with our previously reported model (23) in which Na^+^ is released from the Na2 site when LeuT transitions to a partially inward-open state (IO^2^). To test directly whether the reduction of the Na^+^ affinity in LeuT-F259W is caused by disrupting the integrity of the Na2 site, we made structure-guided mutations that impair Na^+^ binding in the Na1 (N27A) or Na2 (T354A) sites (29). Disrupting the Na2 site in the F259W background (LeuT-F259W/T354A) had only a subtle effect on the Na^+^ binding affinity. In contrast, disrupting the Na1 site in the F259W background (LeuT-N27A/F259W) completely blocked Na^+^ binding (**Fig. 4A**). These results suggest that Na^+^ binds to LeuT-F259W only in the Na1 site, supporting our model that the IO^2^ state corresponds to a conformation in which Na2 (Na^+^ in the Na2 site) has been released (23). In light of our recent MSM and TCF analysis in hDAT that revealed an allosteric coupling between inward-opening and Na2 release (13, 14, 35), these results suggest that the stabilization of F259 in the parallel state promotes release of Na2 through stabilization of the inward-facing intermediate state.

**Figure 4.**
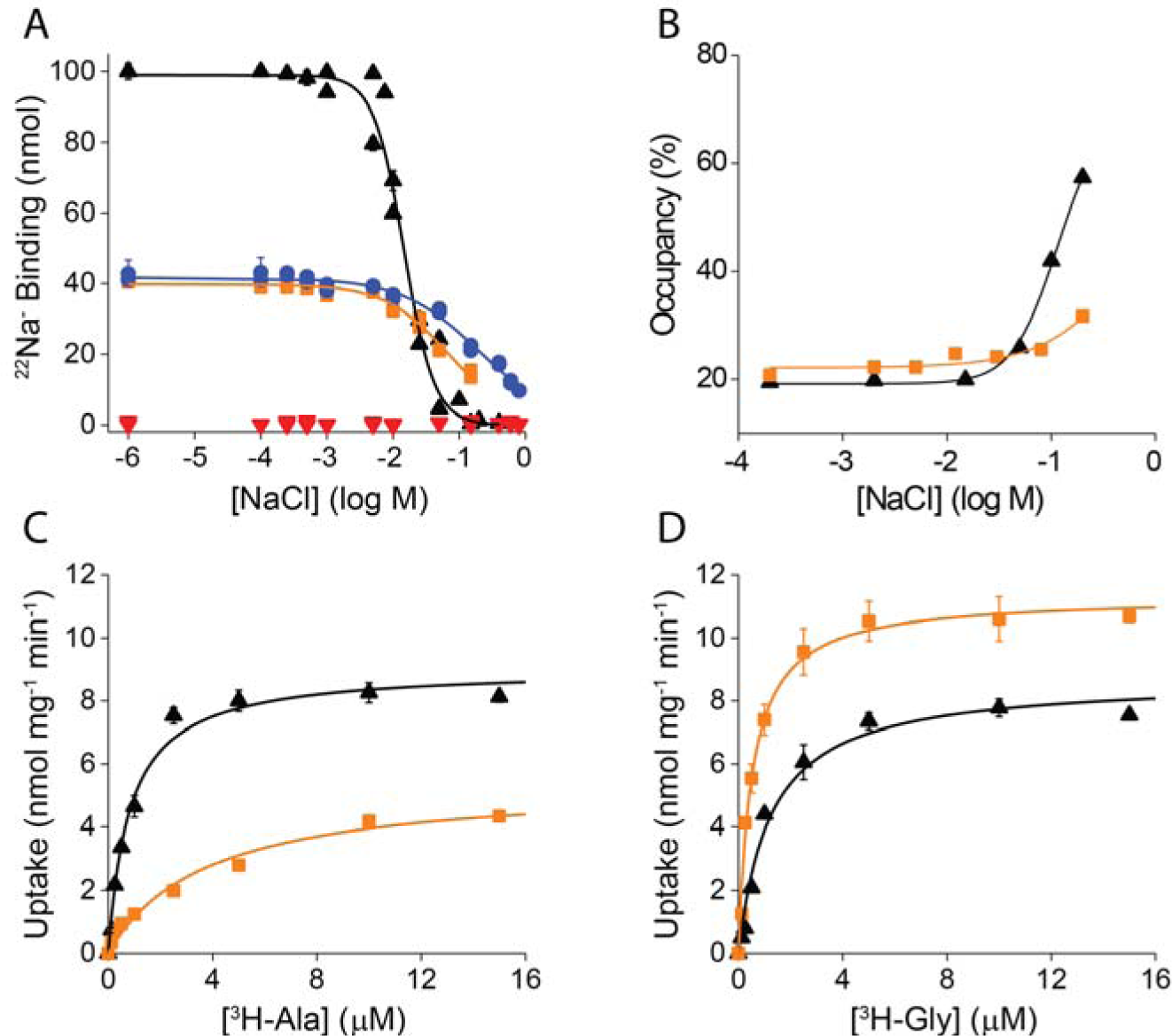
Effect of the F259W mutation on Na^+^ binding and substrate transport. (**A**) Na^+^ binding determined by scintillation proximity assay for LeuT-WT, -F259W, - F259W/T354A, and -N27A/F259W. Binding of 50 μM [^22^Na]Cl (50 Ci/mol) was measured in the presence of increasing concentrations of non-labeled NaCl with 50 ng of the indicated LeuT variant. Data of ≥ 2 independent experiments (with technical triplicates shown as mean ± S.E.M.) were normalized to the binding activity of LeuT-WT and plotted as function of the isotopic displacement of ^22^Na^+^ by non-labeled NaCl. Non-linear regression fitting in SigmaPlot 13 was used to determine the effective concentration of 50 % ^22^Na^+^ displacement (*EC_50_*) and the Hill coefficient for LeuT-WT (black triangle; 14.7 ± 0.9 mM | 1.9 ± 0.2), LeuT-F259W (orange square; 70.3 ± 5.9 mM | 0.9 ± 0.1), LeuT-F259W/T354A (open blue circles; 201.2 ± 17.1 mM | 0.7 ± 0.1), and LeuT-N27A/F259W (red triangles). (**B**) Ensemble averaged occupancy in the high-FRET state from experiments imaging LeuT-WT (black triangles) and -F259W (orange squares) in the absence of substrates and the presence of the indicated concentrations of Na^+^. Lines are fits to dose-response functions with *EC_50_* values of ~110 mM (WT) and >200 mM (F259W). (**C,D**) Uptake of ^3^H-Ala (**C**) or ^3^H-Gly (**D**) by proteoliposomes containing LeuT-WT or -F259W. Uptake was performed in Tris-Mes, pH 8.5 containing 50 mM (LeuT-WT) or 800 mM (-F259W) NaCl (equimolarly replaced with Tris/Mes) using proteoliposomes with an internal pH of 6.5(17). The individual uptake data of 2 independent experiments (with technical triplicates) were averaged and shown as mean ± S.E.M. and plotted as function of the substrate concentration. Data were subjected to non-linear regression fitting in SigmaPlot 13 to calculate the *V_max_* and *K_m_* of LeuT-WT for Ala (9.0 ± 0.3 nmol x mg^-1^ x min^-1^ | 0.8 ± 0.1 µM) and Gly (8.7 ± 0.4 nmol x mg^-1^ x min^-1^ | 1.2 ± 0.2 µM) and of LeuT-F259W for Ala (5.4 ± 0.5 nmol x mg^-1^ x min^-1^ | 3.7 ± 0.9 µM) and Gly (11.3 ± 0.3 nmol x mg^-1^ x min^-1^ | 0.5 ± 0.06 µM).

### The F259W mutation increases selectivity for Gly transport

Previously it has been observed that the size/volume of the amino acid side chain of a substrate is correlated with the substrate’s transport efficiency (19, 45), including a four-fold decrease in transport rates for Gly relative to Ala, a result that is not predicted by our model. To assess the functional relevance of a high basal occupancy of the IO^2^ state, we compared the transport rates of radiolabeled Gly and Ala by LeuT-WT and the F259W mutant. In contrast to previous findings, and in accordance with our model, we find that, in fact, Gly and Ala are transported by WT LeuT with nearly equal maximal velocity (*V_max_*)(**Table S1**). We note that the previous uptake experiments (19, 45) were performed in the absence of a pH gradient across the membrane of the LeuT-containing proteoliposomes (i.e., pH_in_ < pH_out_). Hence, these investigations lacked a critical component needed for optimal uptake measurements for LeuT and other Cl^-^-independent bacterial NSS homologues that feature H^+^ extrusion as part of the Na^+^/substrate symport reaction(46–49). We therefore conclude that these experimental differences account for the discrepancy in the transport rates. In addition, the F259W mutation was found to increase the *K_m_* and decrease the *V_max_* of Ala transport, leading to substantially diminished transport activity for this prototypical LeuT substrate (**Fig. 4C**). This observation is consistent with the idea that the increased steric occupancy of the binding site by W259, in combination with the Ala substrate, mimics that observed for the larger substrates in the wild-type LeuT. In contrast, the *K_m_* for Gly was decreased substantially and the *V_max_* was increased compared to WT (**Fig. 4D**). Thus, the F259W mutation is sufficient to increase the selectivity of LeuT for Gly transport. These results are also consistent with recently published results in which the reverse mutation in GlyT (i.e., Trp-to-Phe at the position corresponding to F259 in LeuT) transformed the normally selective transporter into a non-selective amino acid transporter (50).

## Discussion

We demonstrate here a mechanism by which substrate selectivity is achieved in the function of the NSS homologue LeuT. Our previous NbIT analysis (24) predicted that residues F259 and I359 serve as the main mediators of allosteric communication between the substrate in the S1 site and the intracellular gate. Here, we used a combination of MD simulations, QM/MM calculations, smFRET imaging, and Na^+^ binding assays to test the prediction and find that the mode of interaction of substrates with these residues elicits substrate-specific allosteric modulation of both intracellular gating and Na^+^ release from the Na2 site. In this allosteric mechanism, F259 acts in a manner analogous to a volumetric sensor of substrate bulk. Low-volume substrates that bind while allowing free rotation of F259 induce rapid sampling of an inward-facing, intermediate FRET state associated with the release of Na^+^ from the Na2 binding site (IO^2^). In contrast, bulky substrates that act sterically to prevent the rotation of F259 stabilize the fully Na^+^-bound inward-closed state. We show that the F259W mutation fixes the residue in the parallel rotameric state and induces rapid sampling and high occupancy of the inward-facing intermediate FRET (IO^2^) state. In the IO^2^ state, the affinity of Na^+^ in the Na2 site is drastically reduced even in the absence of substrate. Together, these results suggest that in the case of F259, occupancy of the parallel state is necessary and sufficient to achieve the inward-facing, intermediate FRET state (IO^2^) associated with rapid substrate transport (23).

We note, however, that the relationship between apparent intermediate state occupancy and transport rates (*V_max_*) is not linear. Thus, Val is transported much more rapidly than Leu, but partially stabilizes the inward-closed state while destabilizing the parallel F259 rotamer, albeit to a lesser extent than Leu. One possible explanation for this non-linearity is that only brief transitions to the inward-facing, intermediate FRET state are required, such that the rapid exchange between perpendicular F259 states observed in Val are able to facilitate brief transitions to the IO^2^ state, which are difficult to resolve at our present experimental time resolution.

The smFRET imaging results described here gain additional perspective from recent site-directed fluorescence quenching spectroscopy (SDFQS) experiments (45) that provided evidence for substrate-induced conformational changes. Specifically, an increase in solvent accessibility of a fluorophore inserted at the intracellular end of TM5 was observed upon binding of all transported substrates, but not binding of inhibitors or Na^+^ alone. With the labeling pattern used in the smFRET experiments, such a conformational change would not be detected, and would instead be observed as intracellular gate closing. The SDFQS experiments showed a substrate-dependent decrease in fluorophore solvent accessibility correlated with substrate size and transport rates. The rank order of these effects was the same as observed here for the F259 rotamer dynamics and the related inward-facing, intermediate-FRET state stabilization and increased intracellular dynamics. Thus, the modulation in accessibility observed with SDFQS may arise from the increased inward-facing, intermediate-FRET state occupancy observed with the more rapidly transported substrates, or to a mechanistically related process.

In view of the determinant role of F259 interactions with substrates demonstrated here, the conservation of F259 across the synaptic monoamine transporters (sMATs) becomes of interest. The dopamine, norepinephrine, and serotonin transporters (DAT, NET, and SERT, respectively) all transport substrates with bulky ring substituents. Thus, it may be expected that the residue homologous to F259 in each of these transporters would engage these substrates in a manner that would inhibit transport. While comparison of X-ray structures of *Drosophila* DAT (dDAT) (51–53) in the presence of various S1 ligands indicate that the corresponding F325 can sample distinct states similar to the parallel and perpendicular states observed in our MD simulations of LeuT, extant structure data do not reveal a clear correlation between the F325 rotamer state and the bound ligands in dDAT. In dDAT crystal structures, which are all in either an occluded or outward-facing open state, the F325 residue is in a perpendicular state when interacting with either the endogenous substrate dopamine or the inhibitor cocaine (53), while nisoxetine, a selective NET inhibitor in humans that also binds to and inhibits dDAT, contacts F325 with one of its rings to stabilize a parallel-like state of F325(51). Regarding the possibility that crystallographic conditions could be responsible for the stabilization of F325 in a particular state in the dopamine-bound structure, similar to what we reasoned above based on the QM/MM calculations on the LeuT X-ray structures, we note that significant occupancy of the parallel state was also not seen in our extensive MD simulations of inward-opening in dDAT and the human homologue hDAT(13, 14, 35). This observation suggests that in DAT, dopamine does in fact stabilize the perpendicular state regardless of intracellular gating configuration. This inference is consistent with the transfer of the regulation of inward opening and Na2 release in eukaryotic transporters to the intracellular N-terminus, as described recently (54). In X-ray structures of human SERT (hSERT) (55, 56) bound to the antidepressant inhibitors paroxetine and (S)-citalopram (55) the corresponding F341 is significantly displaced from its position in LeuT and dDAT, as it is situated below the inhibitors with the ring in a perpendicular-like state, while hSERT in complex with sertraline and fluvoxamine (56), F341 exhibits a parallel-like state. Notably, no structure of SERT with substrate has been published to date, perhaps reflecting the dynamic nature of this state anticipated by our smFRET measurements of LeuT.

The influence of inhibitors on F341’s χ1 angle has recently been associated with the affinity of inhibitors (57). The diverse conformational space sampled by this conserved phenylalanine in published sMAT X-ray structures may indicate that the mechanistic role it plays in substrate transport in LeuT is not strictly conserved across the sMATs but has evolved to require an interplay with other structural elements (54). In view of its conservation, however, the predicted role of this residue in the function of this important transporter family warrants further investigation, as the design of inhibitors that specifically act via the inhibition of the Na^+^ release step could lead to the development of novel antidepressants and psychostimulants.

## Acknowledgements

We thank Roger Altman for preparation of microfluidic chambers and the members of the Blanchard, Javitch, and Weinstein labs for helpful discussions. This research was supported by NIH grants R21 MH099491 (S.C.B.), U54 GM087510 (S.C.B., J.A.J., H.W.), P01 DA012408 (H.W.), R01 DA041510 (J.A.J., H.W., S.C.B.), and a Ruth L. Kirschstein National Research Service Award F31 DA035533 to M.V.L. Computational results utilized in this work were carried out at resources of the Oak Ridge Leadership Computing Facility (ALCC allocation BIP109) at the Oak Ridge National Laboratory, which is supported by the Office of Science of the U.S. Department of Energy under Contract No. DE-AC05-00OR22725; an allocation at the National Energy Research Scientific Computing Center (NERSC, repository m1710) supported by the Office of Science of the U.S. Department of Energy under Contract No. DE-AC02-05CH11231; the Extreme Science and Engineering Discovery Environment (XSEDE, account TG-MCB120008), which is supported by National Science Foundation grant number ACI-1053575; and the Anton 2 supercomputer provided by the Pittsburgh Supercomputing Center (PSC) through Grant *R01GM116961* from the National Institutes of Health. The Anton 2 machine at PSC was generously made available by D.E. Shaw Research. The computational resources of the David A. Cofrin Center for Biomedical Information in the HRH Prince Alwaleed Bin Talal Bin Abdulaziz Alsaud Institute for Computational Biomedicine at Weill Cornell Medicine are gratefully acknowledged.

## Author contributions

M.V.L and G.K. performed the MD simulations and M.V.L. analyzed the data. M.V.L and Z.S. performed the QM/MM calculations and Z.S. analyzed the data. D.S.T. performed single molecule FRET experiments and analyzed the data. M.Q. performed radiolabeled substrate binding and transport assays. All authors contributed to experimental design, interpretation of data, and writing the manuscript.

## Competing financial interests

S.C.B. has an equity interest in Lumidyne Technologies.

## Code Availability

The full source code of SPARTAN(32), which was used for all analysis of smFRET data, is publicly available: http://www.scottcblanchardlab.com/software

## Data Availability

The data that support the findings of this study are available from the corresponding authors upon reasonable request.

## Methods

### Molecular Constructs

The X-ray structure of LeuT from PDBID 3F3E (19) was used for all atomistic MD simulations. In this structure, LeuT is bound to Leu ligand and two Na^+^ ions in the Na1 and Na2 binding sites. Short loop fragments (residues 1-4 and 132-134), not resolved crystallographically, were built and added to the X-ray structure using Modeller (58). To build Ala-, Val-, and Gly-bound LeuT models, ligand Leu in the 3F3E structure was substituted by the respective ligands using the Mutator plug-in within VMD (59) (the two Na^+^ ions were retained). In this manner, starting protein structures in complex with these different ligands for subsequent MD simulations were identical.

Using CHARMM-GUI webserver (60), the LeuT models were embedded in a membrane with 75:25 mixture of POPE (1-palmitoyl-2-oleoyl-sn-glycero-3-phosphoethanolamine) / POPG (1-palmitoyl-2-oleoyl-*sn*-glycero-3-phospho-(1'-*rac*-glycerol)) lipids (296 lipids in total), hydrated and ionized with 0.15 M Na^+^Cl^-^ salt, resulting in the final system size of ~118,000 atoms (including hydrogen atoms).

### Molecular Dynamics (MD) Simulations

All the molecular systems assembled were first subjected to a multi-step equilibration protocol (for examples see (13, 35)) with NAMD version 2.9 using CHARMM36 parameters for proteins (61), lipids (62), and ions. Briefly, this phase included: 1) minimization for 5,000 steps and running MD with 1 fs integration time-step for 250 ps, fixing all atoms in the system except for the lipid tails; 2) minimization for 2,500 steps and performing MD with 1 fs time-step for 500 ps with constrained protein backbone and lipid headgroups (force constant of 1 kcal/(mol Å^2^) and keeping water out of the membrane hydrophobic core; 3) gradual release of the constraints on the protein backbone and lipid headgroup atoms (force constant of 0.5 and 0.1 kcal/(mol Å^2^)) while still keeping water out of the membrane interior. At each value of the force constant, the system was minimized for 2,500 steps followed by 500 ps MD (with 1 fs time-step); 4) unbiased MD simulation for 30 ns using 2 fs time-step. These steps implemented PME for electrostatics interactions (63), and were carried out in the NPT ensemble under semi-isotropic pressure coupling conditions, at 310 K temperature. The Nosé-Hoover Langevin piston (64) algorithm was used to control the target *P*=1 atm pressure with the *LangevinPistonPeriod* set to 100 fs and *LangevinPistonDecay* set to 50 fs. The van der Waals interactions were calculated applying a cutoff distance of 12 Å and switching the potential from 10 Å.

After this initial phase, the molecular systems were subjected to 3 microsecond long MD simulations on Anton1, a special-purpose supercomputer machine (65). These production runs implemented the same set of CHARMM36 force-field parameters and were carried out in the NPT ensemble under semi-isotropic pressure coupling conditions (using the Multigrator scheme that employs the Martyna-Tuckerman-Klein (MTK) barostat (66) and the Nosé-Hoover thermostat (67)), at 310 K temperature, with 2 fs time-step, and using PME for electrostatic interactions. All the other run parameters were derived from the Anton guesser scripts based on the system chemistry.

A representative frame from the Anton trajectory of Gly-bound LeuT in which F259 ring assumed “parallel state” (see Results) was chosen to mutate the F259 residue into Trp (F259W) using the Mutator plug-in in VMD. The resulting molecular system was minimized and ran with unbiased MD for 30 ns using NAMD 2.9 after which it was transferred to Anton1 for 3 microsecond MD simulation.

### Hybrid Quantum Mechanics / Molecular Mechanics (QM/MM) Calculations

Hybrid quantum mechanics/molecular mechanics (QM/MM) calculations scanning the angle spaces of F259 and I359 were performed using the QSite program from version 2017-2 of Maestro from Schrödinger (68). In order to maintain consistency with the MD simulations, the protein models were prepared from the leucine-bound crystal structure (PDB ID 3F3E) (19). Structures for the glycine-, alanine-, and valine-bound LeuT were prepared by mutating the bound leucine in Maestro. The QM portion of the calculations were performed using the density functional theory method DFT-B3LYP (69, 70) with the LACVP* basis set. The QM region in all cases included the bound ligand, F259, I359, and the side chain of the scanned residue’s two immediate neighbors in sequence. (G258, F259, F260, and the side chain of I359 when F259 was scanned; A358, I359, M360, and the side chain of F259 when I359 was scanned). The MM region was comprised of the rest of the protein, with the backbone atoms constrained using a 25.00 kcal/(mol Å^2^) harmonic force constant. Convergence of MM minimization steps was based on an energy change criterion of 10^−7^ kcal/mol and a gradient change criterion of 0.01 kcal/(mol•Å), using a Truncated Newton algorithm, and a maximum of 1000 cycles. After an initial minimization step, the dihedral being scanned (F259 χ₂ and I359 χ₁) was rotated a total 1080° in steps of 5°. The initial dihedral was based on that of the crystal structure (81.612° for F259 χ₂, 226.799° for I359 χ₁).

### Identification of the F259 Rotameric States

In order to characterize the F259 rotameric state, we calculated the dihedral angle formed by the C_α_, C_β_, C_γ_, and C_D1_ carbons across each MD trajectory. As we introduced new substrates into the S1 binding sites, the initial dynamics of the binding site residues are expected to be far from equilibrium. We used a recently developed method (71), adapted for angular data, to determine how much of the initial portion of the trajectories should be discarded as equilibration phase. In the analysis, the first *t_0_* frames are discarded, where *t_0_* is chosen such that the effective number of observations remaining after correcting for autocorrelation is maximized. Given a trajectory of time length *T*, we calculate the effective number of observations in the time window from *t_0_* to *T*, where *t_0_* ranges from 0 to *T*. The effective number of observations was calculated as:

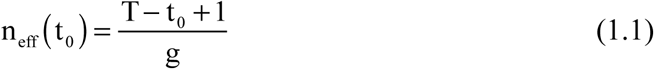

where g is the statistical efficiency:

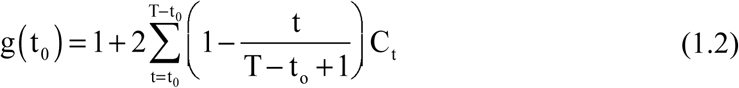

and *C_t_* is the time-lagged circular correlation,

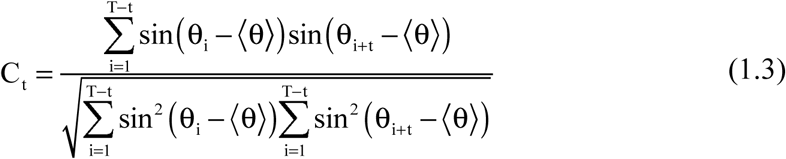

After removing the non-equilibrium portions of the trajectories (**Fig. S1-5**), we performed k-centroids clustering using the kcca function in the flexclust package in R (72) to identify the rotameric states. The angular histograms of the glycine- and alanine-bound simulations revealed 4 distinct states, and thus the number of clusters, *k*, was chosen to be 4 and clustering was performed using a modified k-means algorithm that minimizes the angle between cluster members and the standardized mean of the cluster. We clustered the angle distributions from the glycine-bound trajectory, which sampled all 4 states equally, and then used the resulting centroids and the predict function to cluster the alanine-, valine-, leucine-, and F259W glycine-bound angle distributions. Finally, the symmetric clusters were merged for each distribution.

### Protein expression and purification for smFRET experiments

LeuT constructs were expressed and purified as described previously (23). Briefly, LeuT was expressed in *E. coli* using pQO18-TEV vector derivatives with the H7C/R86C mutations or H7C/R86C/F259W as previously described(73). The constructs were solubilized in n-dodecyl β-D-maltopyranoside (DDM), purified by immobilized metal affinity chelate chromatography (IMAC) using a Ni^2+^ Sepharose 6 FastFlow column (GE Healthcare), and then labeled with maleimide-activated LD550 and LD650 fluorophores (Lumidyne Technologies) at a 1:1.5 molar ratio (200 µM total) for 1 hr at 4 °C, followed by size exclusion chromatography using a Superdex 200 16/60 column.

### smFRET imaging experiments

smFRET imagine experiments were performed using a prism-based total internal reflection fluorescence (TIRF) microscope as previously described (21–23, 32). Passivated microfluidic imaging chambers were prepared with 0.8 µM streptavidin (Invitrogen) and 4 nM biotin-*tris*-(NTA-Ni^2+^)(74) and fluorophore labeled, His-tagged LeuT molecules were reversibly surface-immobilized. LD550 fluorophores were excited by the evanescent wave generated by total internal reflection (TIR) of a single-frequency light source (Opus 532, Laser Quantum). Photons emitted from LD550 and LD650 were collected using a 1.27 NA 60× water-immersion objective (Nikon) and a MultiCam-LS device (Cairn) with a T635lpxr-UF2 dichroic mirror to separate the spectral channels onto two synchronized sCMOS cameras (Flash 4.0 v2, Hamamatsu). Fluorescence data were acquired at 10 frames per second (100 ms time resolution) using custom software implemented in LabView (National Instruments).

All experiments were performed in buffer containing 50 mM Tris/Mes (pH 7.0), 10% glycerol, 0.02% (w/v) DDM (Anagrade, Anatrace), 1 mM 2-mercaptoethanol and 200 mM total salt (NaCl and KCl, as specified). An oxygen-scavenging environment containing 0.2 unit per mL glucose oxidase (Sigma G2133), 1.8 units per μL catalase (Sigma C40), 0.1% (v/v) glucose was employed to minimize photobleaching. Both enzymes were purified by gel filtration using a Superdex 200 13/30 column (GE Healthcare) prior to use. All experiments were performed at 25 °C.

Analysis of smFRET data was performed using SPARTAN, freely available smFRET analysis software written in MATLAB (MathWorks) (32), as previously described (23). Spectral bleed-through from the donor to the acceptor channel was corrected by subtracting a set fraction (0.165) of the donor intensity from the acceptor. The FRET efficiency was calculated as: *E_FRET_ = I_A_/(I_A_+I_D_)*, where *I_A_* and *I_D_* are the donor and acceptor fluorescence traces and *E_FRET_* is set to zero whenever the donor was in the dark state. A subset of the acquired traces was selected for further analysis using the following criteria: (1) single-step donor photobleaching, (2) SNR_Background_ ≥ 15, (3) SNR_Signal_ ≥ 4, (4) <4 donor blinking events, and (5) FRET efficiency above 0.15 for at least 300 frames (30 seconds). Replicates (*n*) are defined as data acquired from independent immobilizations of LeuT, generally performed on separate days with newly prepared buffer solutions and frozen aliquots obtained from a single preparation. Finally, traces were fit to a three-state model using the segmental K-means algorithm (75) as described previously (23).

### Radiotracer-based binding and transport studies

Plasmid-encoded LeuT variants were produced in *E. coli* C43(DE3) and purified as described previously (17). Direct binding of 50 μM [^22^Na^+^]Cl (50 Ci/mol) by 50 ng of purified and desalted protein was measured with the scintillation proximity assay (SPA) as described (17, 76) in assay buffer composed of 100-900 mM Tris/Mes, pH 7.5/0-800 mM NaCl (equimolar substitution of NaCl with Tris/Mes)/20% glycerol/0.1 mM TCEP/0.1% DDM using 1.25 mg/mL copper His-tag PVT SPA beads (Perkin Elmer). Total binding was corrected for the non-proximity-based signal to determine the specific binding activity of each LeuT variant and data were normalized to the maximal binding observed for LeuT-WT that was assayed in parallel in all individual measurements. Data of ≥ 2 independent experiments shown as mean ± SEM of triplicate determinations were subjected to global fitting in SigmaPlot 13 to determine the kinetic constants (the error indicates the SEM of the fit).

Transport of [^3^H]Ala (56 Ci/mmol) or [^3^H]Gly (60 Ci/mmol) or [^3^H]valine (60 Ci/mmol), or [^3^H]Leu (100 Ci/mmol) (all from American Radiolabeled Chemicals, Inc.) was measured in proteoliposomes containing the indicated LeuT variant that was incorporated in pre-formed liposomes composed of *E. coli* polar lipids and POPC (3:1 w/w) at a protein-to-lipid ratio of 1:150 (w/w) as described (17). Uptake of the radiolabeled amino acid was performed in 850 mM Tris/Mes, pH 8.5/50 mM NaCl (for LeuT-WT) or 100 mM Tris/Mes, pH 8.5/800 mM NaCl (for LeuT-F259W) and samples were filtered through 0.45 µm membrane filters (EMD Millipore) on a rapid filtration station (17). Data of 2 independent experiments performed in triplicate were plotted and analyzed in SigmaPlot 13 and the Michaelis-Menten transport constant (*K_m_*) and maximum velocity (*V_max_*) were obtained with non-linear regression fitting using the Michaelis-Menten model. The amount of LeuT incorporated into the proteoliposomes was determined using densiometric quantification of samples subjected to 11%-SDS-PAGE followed by silver staining of the proteins using the ImageJ software (NIH). Known amounts of LeuT (determined with the Amidoblack Protein Assay (77)) served as calibration standards.

