## Supplement for "The allosteric mechanism of substrate-specific transport in SLC6 is mediated by a volumetric sensor"

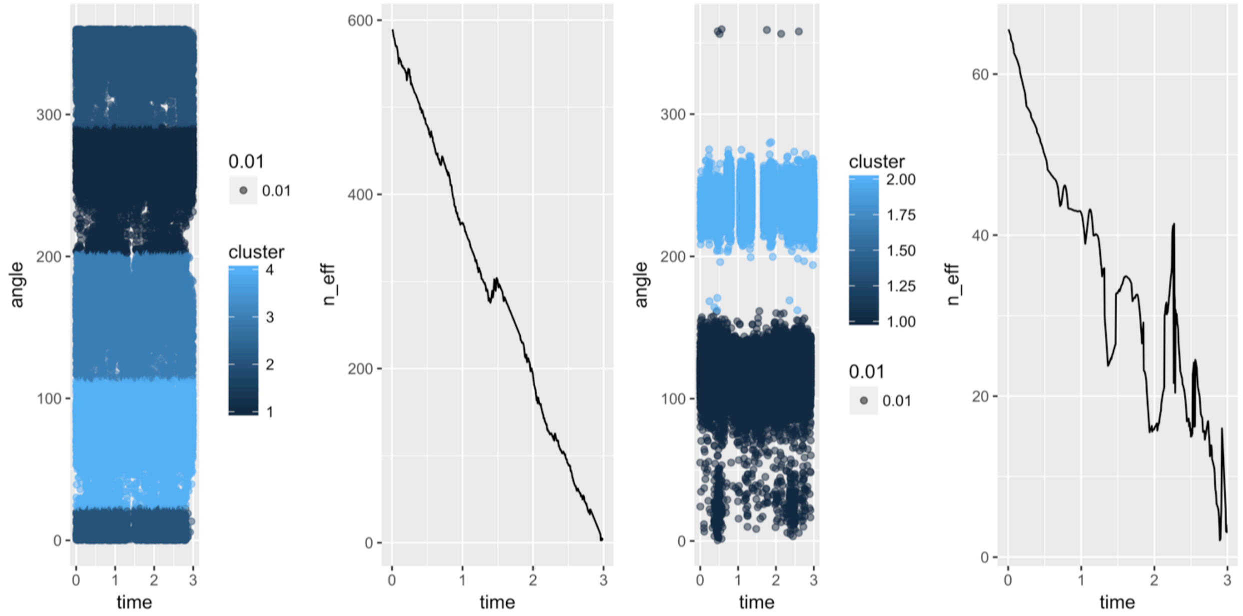


**Figure S1.** Convergence of F259 (left) and I359 (right) in the presence of Gly. For each angle, the time series generated by the trajectory is shown on the left, colored by the state determined by angular k-means clustering of the Gly-bound trajectory, and the estimated effective number of observations as a function of how much of the trajectory is discarded is shown on the right.

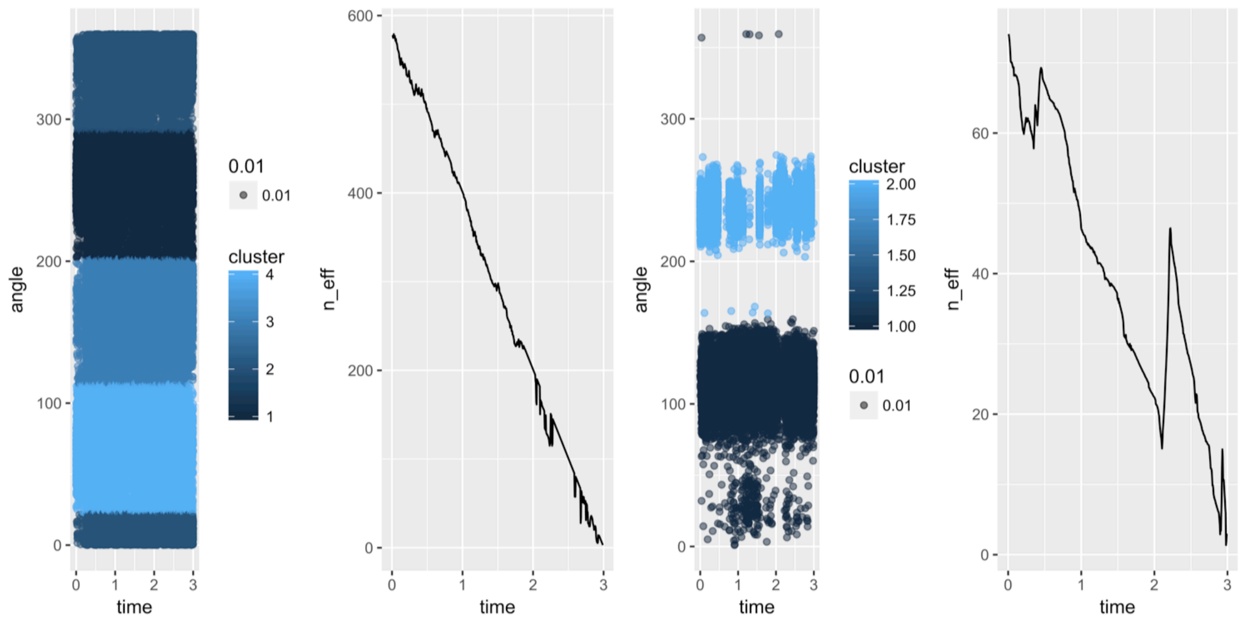


**Figure S2.** Convergence of F259 (left) and I359 (right) in the presence of Ala. For each angle, the time series generated by the trajectory is shown on the left, colored by the state determined by angular k-means clustering of the Gly-bound trajectory, and the estimated effective number of observations as a function of how much of the trajectory is discarded is shown on the right.

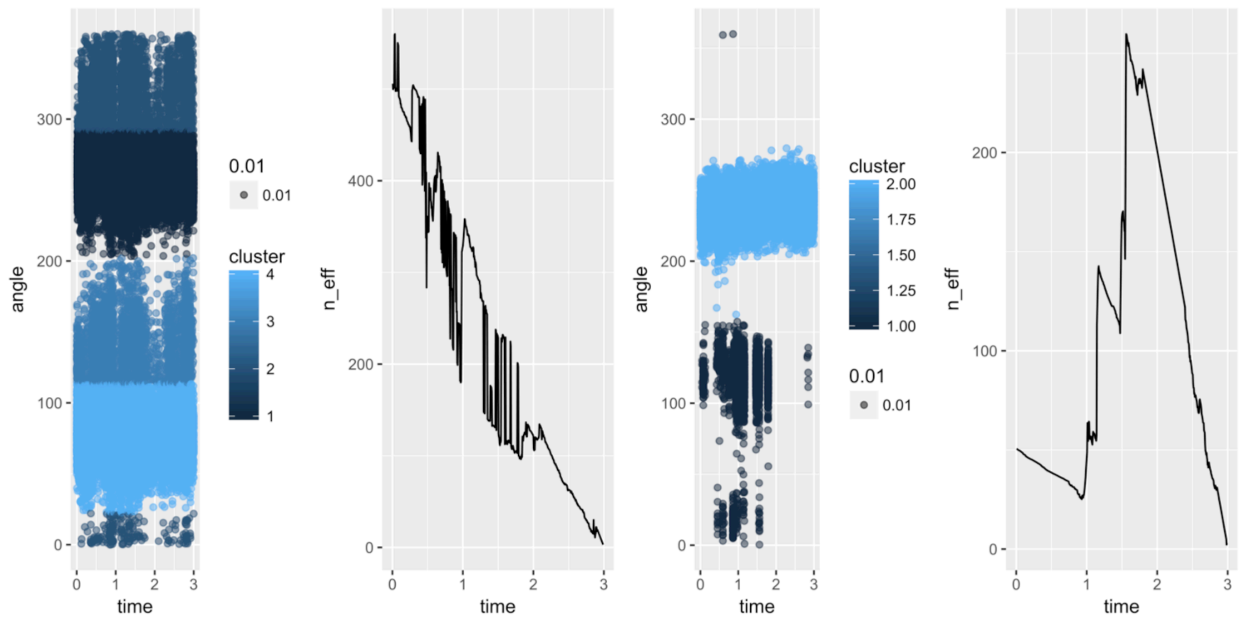


**Figure S3.** Convergence of F259 (left) and I359 (right) in the presence of Val. For each angle, the time series generated by the trajectory is shown on the left, colored by the state determined by angular k-means clustering of the Gly-bound trajectory, and the estimated effective number of observations as a function of how much of the trajectory is discarded is shown on the right.

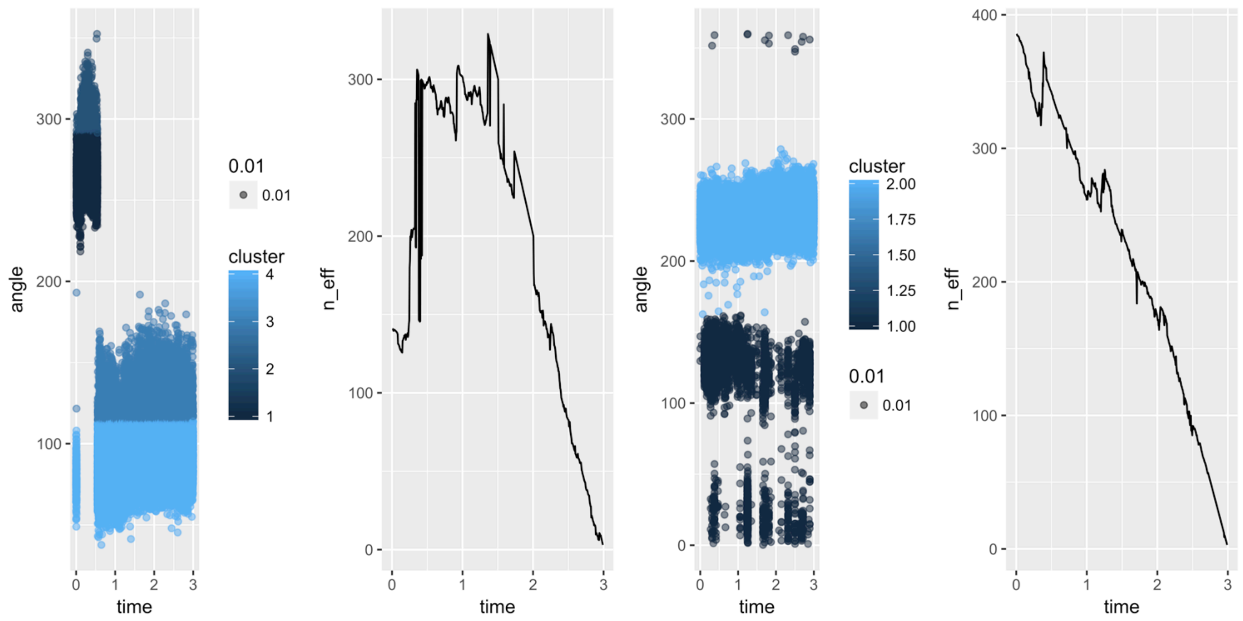


**Figure S4.** Convergence of F259 (left) and I359 (right) in the presence of Leu. For each angle, the time series generated by the trajectory is shown on the left, colored by the state determined by angular kmeans clustering of the Gly-bound trajectory, and the estimated effective number of observations as a function of how much of the trajectory is discarded is shown on the right.

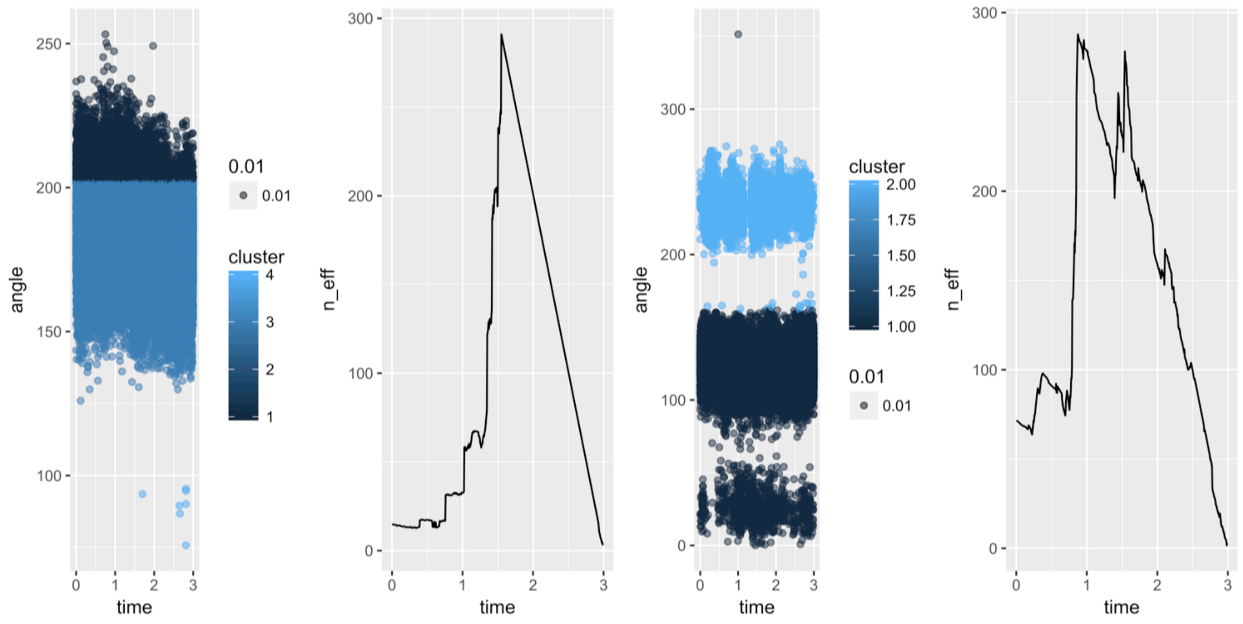


**Figure S5.** Convergence of W259 (left) and I359 (right) in the presence of Gly in the F259W mutant. For each angle, the time series generated by the trajectory is shown on the left, colored by the state determined by angular kmeans clustering of the Gly-bound trajectory, and the estimated effective number of observations as a function of how much of the trajectory is discarded is shown on the right.

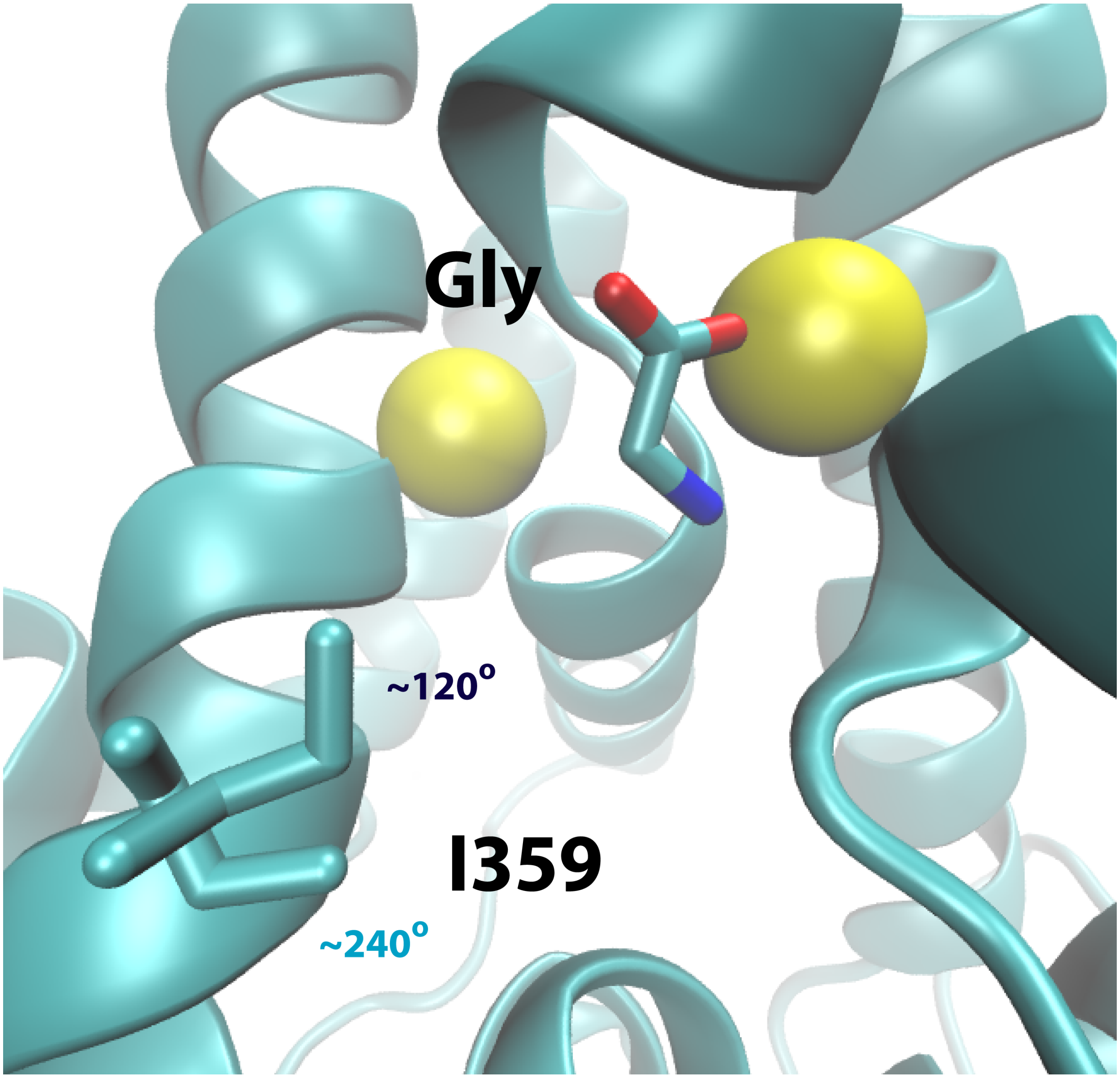


**Figure S6.** Two conformations of I359 in the presence of Gly. The two rotameric states are shown and indicated with the approximate angle of that cluster of states (colored as in Figs. S1-S5, and S8).

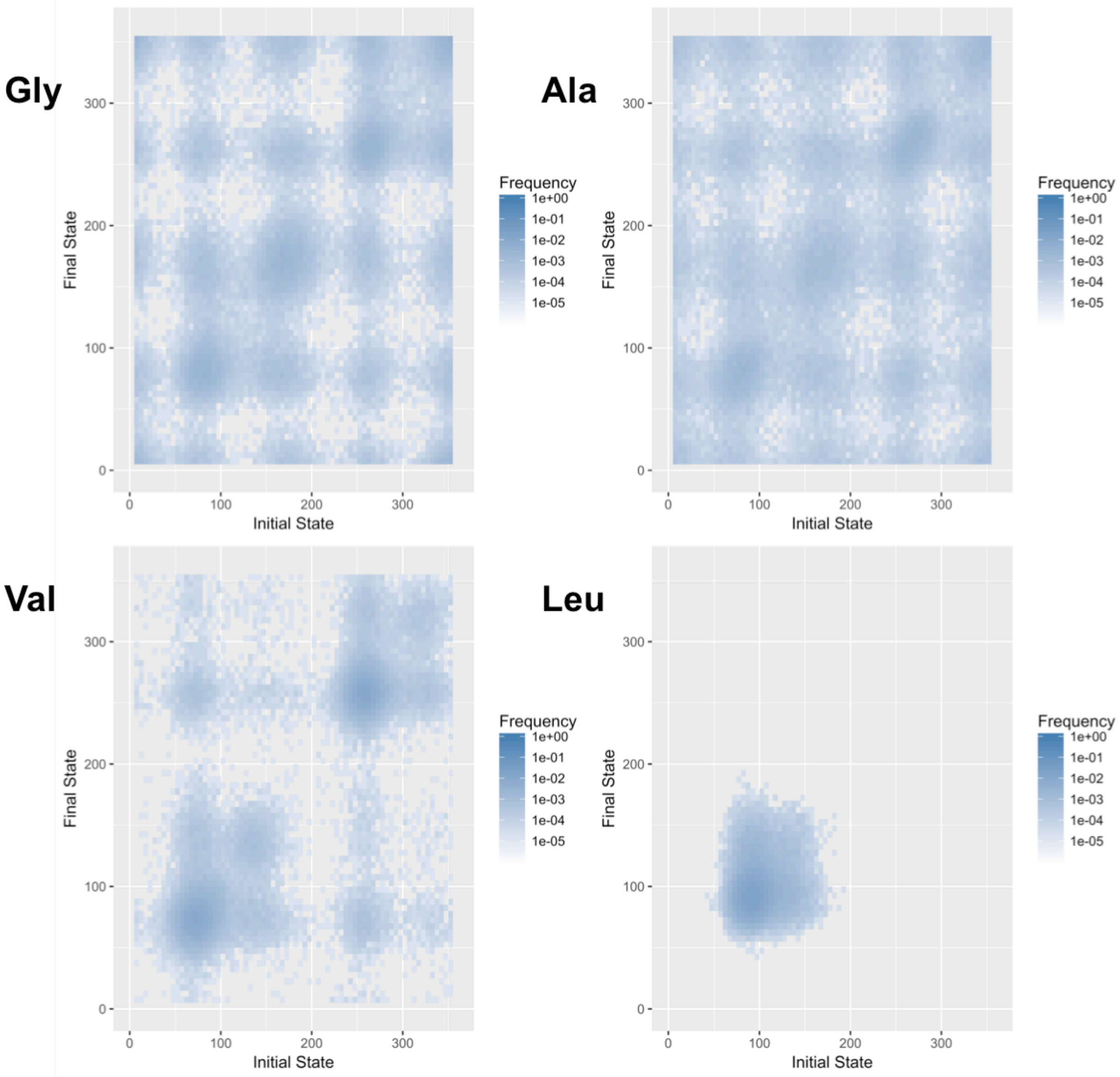


**Figure S7**. Transition density plots for the F259 χ$_2$ angle in the presence of Gly, Ala, Val, and Leu.

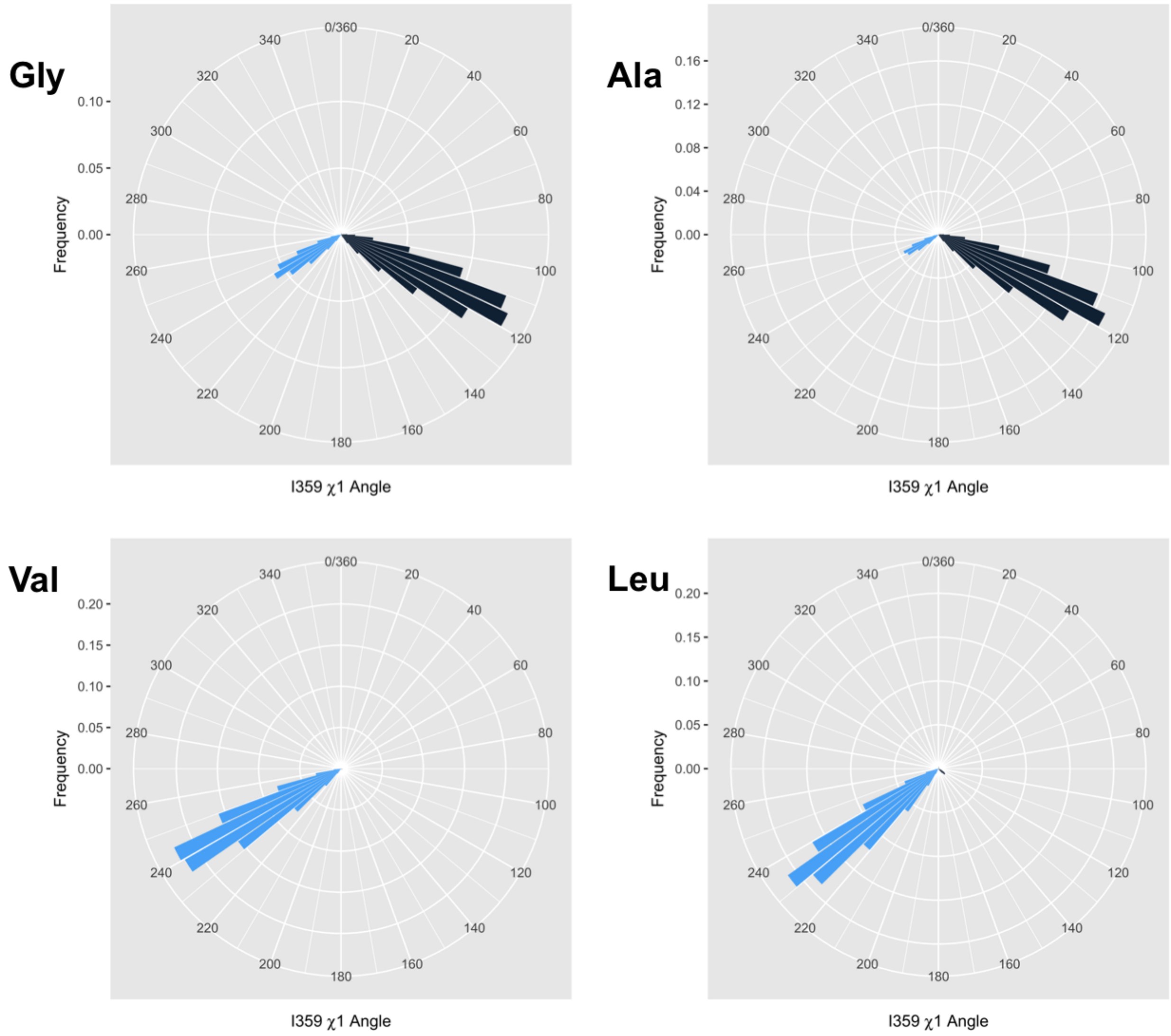


**Figure S8.** Substrates differentially modulate I359 conformation. Rose plots of the angular histograms of the I359 $\chi_2$ angle sampled during the converged portion of the MD simulations are shown, colored by angular kmeans clustering.

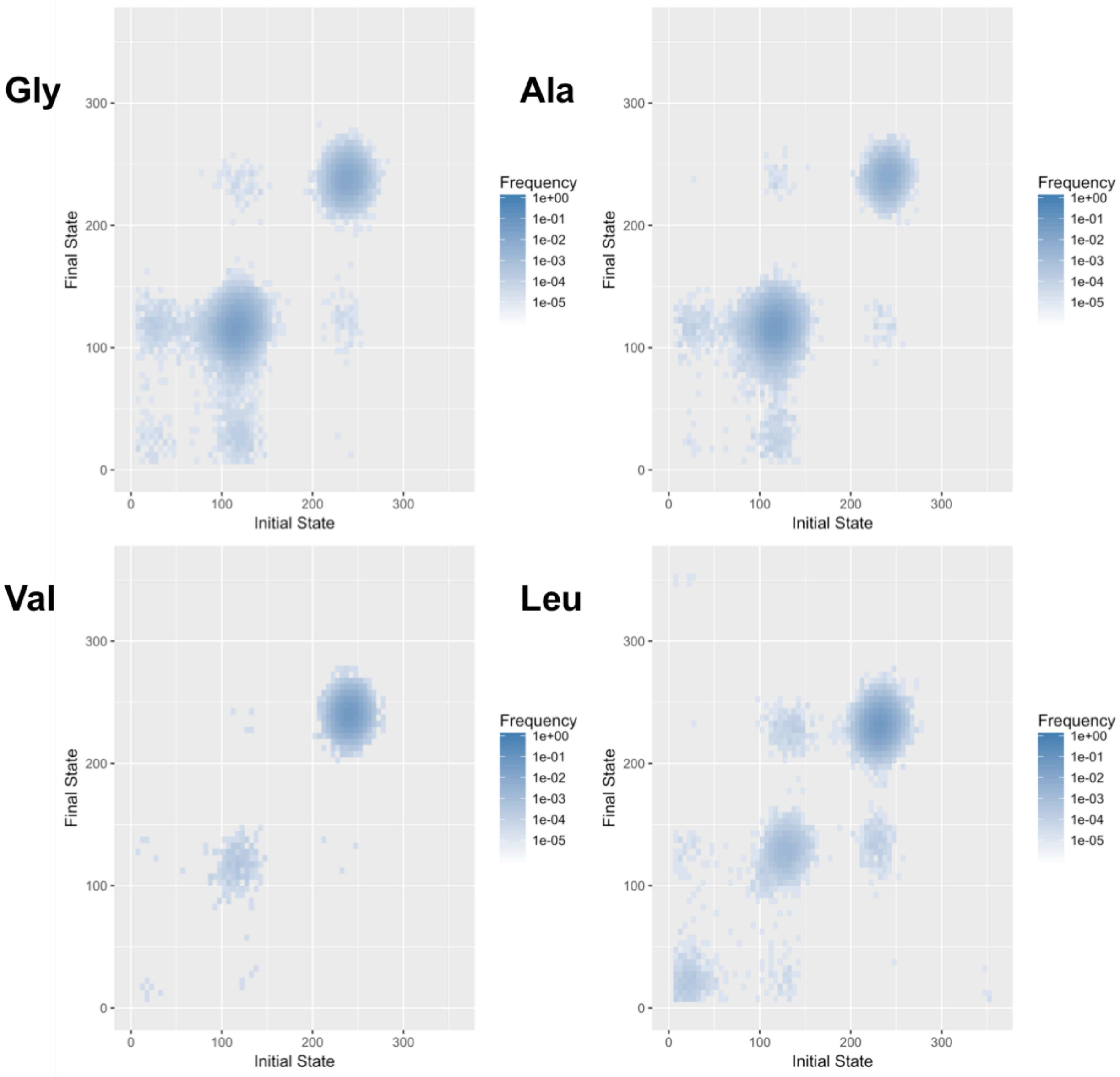


**Figure S9.** Transition density plots for the I359 $\chi_1$ angle in the presence of Gly, Ala, Val, and Leu.

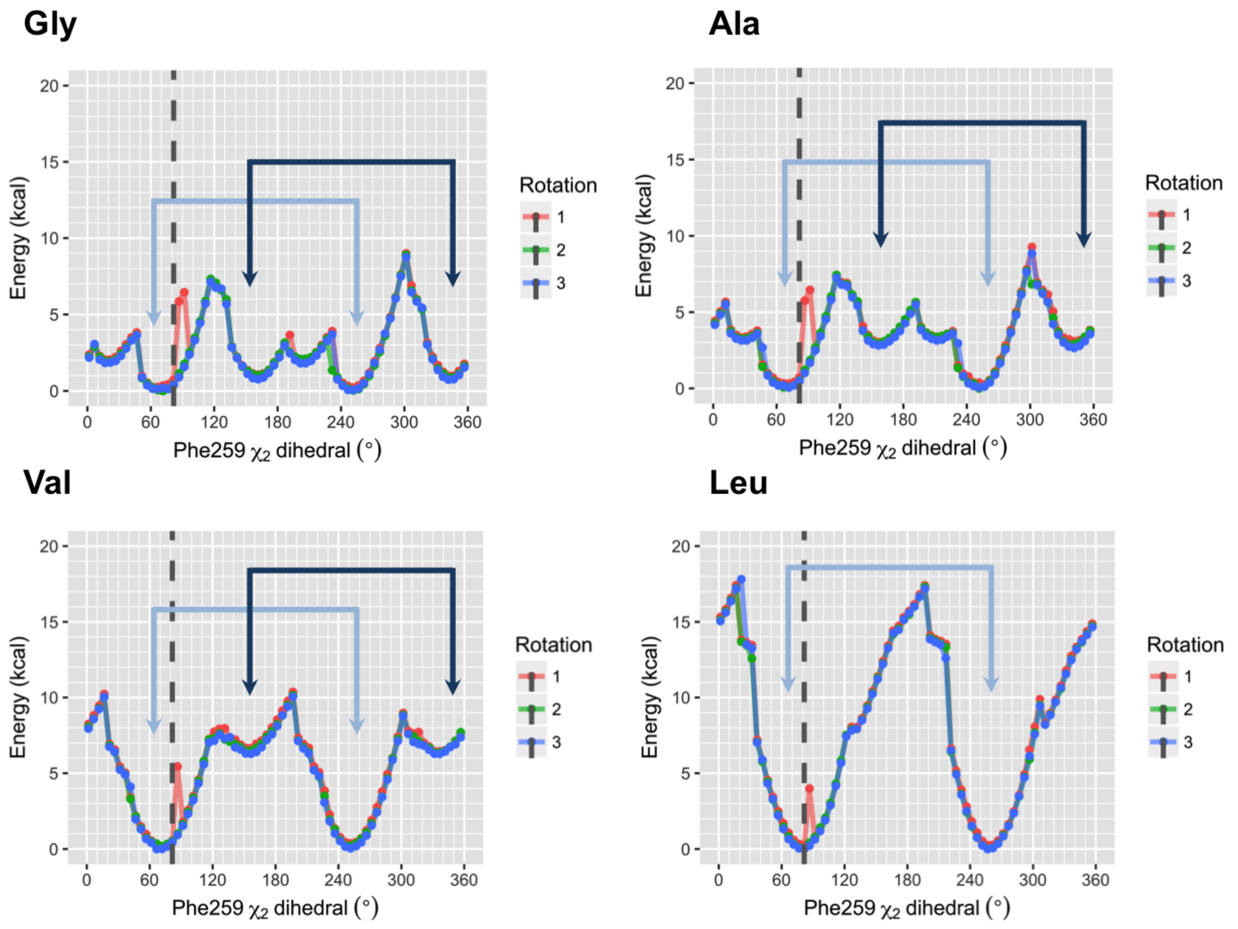


**Figure S10.** QM/MM calculations reveal enthalpically destabilized F259 rotameric states. The F259 $x_2$ dihedral was rotated $5^o$ per step and then the energy of the full system was minimized (see **Methods**). The perpendicular and parallel energy wells are indicated with their respective colors.

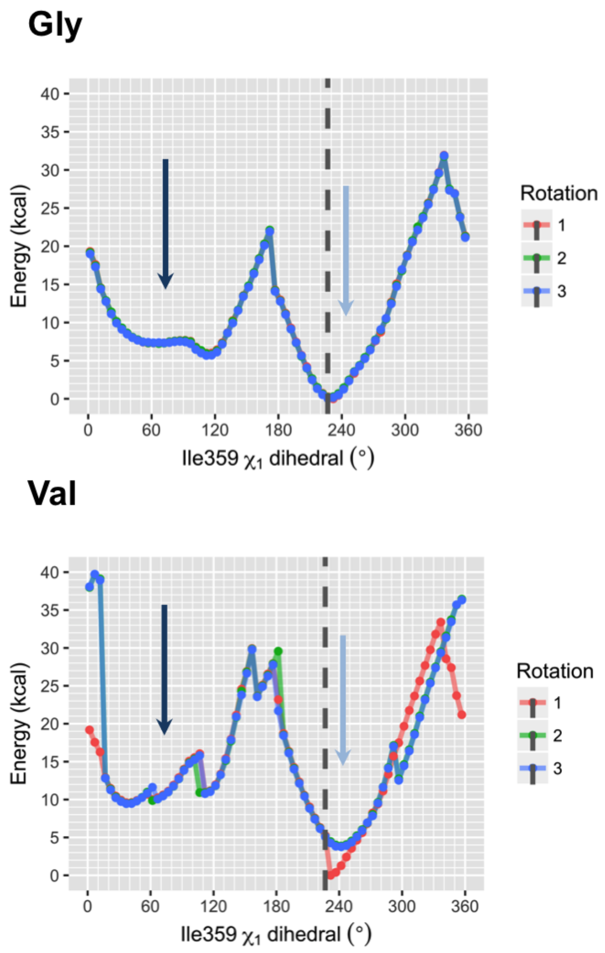


**Figure S11.** QM/MM calculations reveal an enthalpically destabilized I359 rotameric state. The I359 $x_2$ dihedral was rotated 5$^o$ per step and then the energy of the full system was minimized (see **Methods**). The ~120$^o$ and ~240$^o$ wells are indicated with their respective colors.

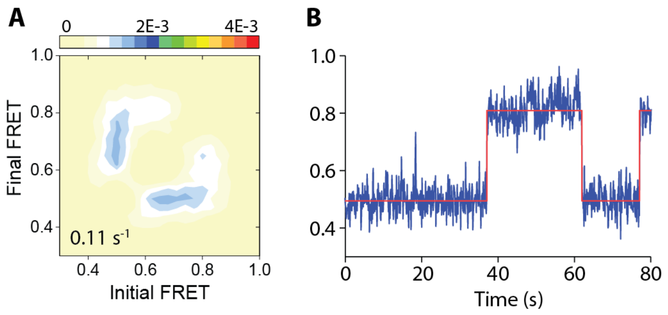


**Figure S12.** Intracellular dynamics of LeuT were imaged in the presence of 30 mM Na+ and in the absence of substrates. Shown are (**A**) Transition density plot and (**B**) and example FRET trace (blue) with state assignment (red). Compare to Fig. 2B-C.

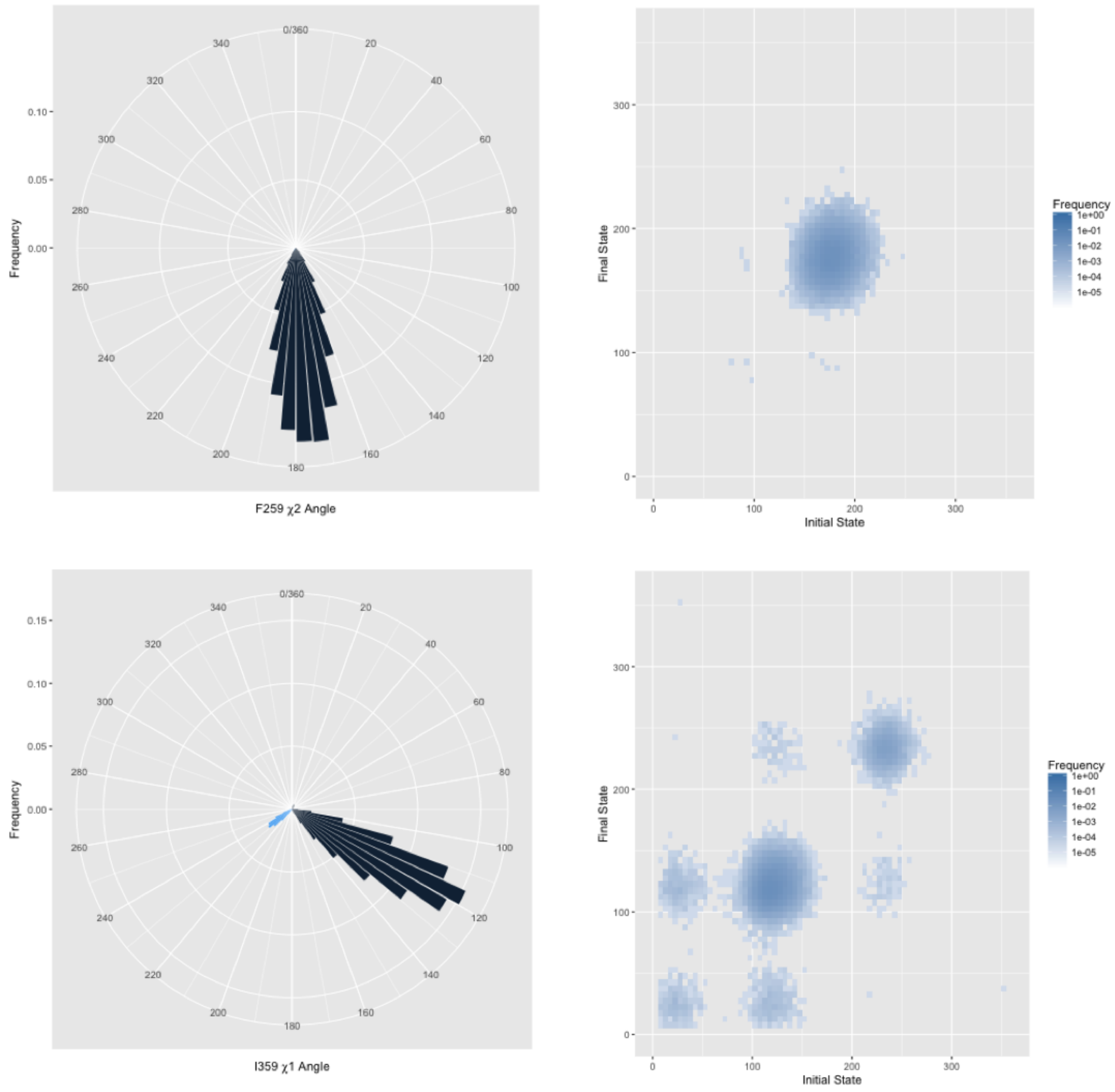


**Figure S13.** W259 and I359 dynamics in F259W. Top: The distribution of W259 $x_2$ angles (left) and the transition density (right). Bottom: The distribution of I359 $x_1$ angles (left) and the transition density (right).

| | | $V_{max}$ (nmol min$^{-1}$ mg LeuT$^{-1}$) | $k_{cat}$ (min$^{-1}$) | $K_M$ (µM) |
|---|---|---|---|---|
| Wild Type | Gly | 8.69 ± 0.40 | 0.50 ± 0.023 | 1.23 ± 0.21 |
| | Ala | 9.03 ± 0.30 | 0.52 ± 0.017 | 0.82 ± 0.11 |
| | Val | 7.67 ± 0.21 | 0.44 ± 0.012 | 0.78 ± 0.89 |
| | Leu | 0.95 ± 0.11 | 0.054 ± 0.006 | 0.41 ± 0.098 |
| F259W | Gly | 11.3 ± 0.29 | 0.65 ± 0.016 | 5.26 ± 0.06 |
| | Ala | 5.40 ± 0.45 | 0.31 ± 0.026 | 3.70 ± 0.85 |

**Table S1**. Kinetic parameters for amino acid transport by LeuT-wild-type and LeuT-F259W. The initial rates of transport of varying concentrations of the indicated amino acids were measured for 30-second periods in the presence of NaCl (50 mM for LeuT-WT and 800 mM for LeuT-F259W). Uptake data were normalized to the actual amount of LeuT incorporated into the proteoliposomes. Densiometric quantification of the intensity of protein bands in proteoliposome samples subjected to SDS-PAGE followed by silver staining of the gel was performed using the ImageJ software (NIH). Data of ≥ 2 independent experiments performed in technical triplicates were subjected to non-linear regression analysis in SigmaPlot 13, and the kinetic constants are shown as the mean ± SEM of the fits.
